# The local and global geometry of trabecular bone

**DOI:** 10.1101/2020.12.02.408377

**Authors:** Sebastien J.P. Callens, Duncan C. Tourolle né Betts, Ralph Müller, Amir A. Zadpoor

## Abstract

The organization and shape of the microstructural elements of trabecular bone govern its physical properties, are implicated in bone disease, and can serve as blueprints for biomaterial design. To devise fundamental structure-property relationships, it is essential to characterize trabecular bone from the perspective of geometry, the mathematical study of shape. Here, we used the micro-computed tomography images of 70 donors at five different sites to characterize the local and global geometry of human trabecular bone, respectively quantified by surface curvatures and Minkowski functionals. We find that curvature density maps provide sensitive shape fingerprints for bone from different sites. Contrary to a common assumption, these curvature maps also show that bone morphology does not approximate a minimal surface but exhibits a much more intricate curvature landscape. At the global (or integral) perspective, our Minkowski analysis illustrates that trabecular bone exhibits other types of anisotropy/ellipticity beyond interfacial orientation, and that anisotropy varies substantially within the trabecular structure. Moreover, we show that the Minkowski functionals unify several traditional morphometric indices. Our geometric approach to trabecular morphometry provides a fundamental language of shape that could be useful for bone failure prediction, understanding geometry-driven tissue growth, and the design of complex tissue engineering scaffolds.

## 1. Introduction

Many natural and man-made materials are characterized by a complex and often hierarchical spatial architecture. A well-known biological example of such a spatially structured material is trabecular bone, exhibiting a characteristic sponge-like morphology [1]. The quantitative morphological characterization of trabecular bone and other structured materials is essential in the study of these systems, for two primary reasons [2]. First, the morphology or architecture of many materials is often the outcome of a biological or physical process. The study of such morphologies, therefore, provides insight into the mechanisms governing their formation. In trabecular bone, for example, the organization of the microstructure is driven by external loading, and changes in the morphology can be indicative of bone diseases, such as osteoporosis [3–5]. Second, the morphology of spatially complex materials can strongly affect their physical properties, making morphological characterization indispensable for establishing structure-function relationships. For example, this intimate structure-property connection enables metamaterials to achieve unique properties [6], and the morphology of biomaterials directly elicits biological responses, affecting aspects such as cell migration, cell fate and spatial tissue organization [7–10].

In the context of trabecular bone, the importance of the microarchitecture has long been recognized, and many morphometric indices have been proposed [11]. However, such indices typically only quantify a particular morphological aspect, such as density, thickness, or interfacial anisotropy, and often lack a fundamental geometric foundation or interpretation [12]. For example, the well-known structure model index (SMI), which classifies trabecular bone by its rod-like or plate-like nature [13, 14], is known to be conceptually flawed by its inability to capture all types of naturally-occurring shapes within the trabecular structure [13, 14]. Moreover, calculating the same metric using different software tools often provides significantly different results, owing to substantial variations in the algorithm implementations [15, 16]. Hence, there is a need for a unifying, robust approach to quantitatively characterize the shape of complex materials, including trabecular bone. Such a well-defined, mathematical framework is established in the realms of differential and integral geometry, providing fundamental descriptors of local and global shape. Local shape can be accurately captured using the concept of surface curvature. For any small neighborhood on a surface, the mean and Gaussian curvatures, defined in terms of the principal curvatures, capture the most fundamental shape information (Figure 1). The magnitudes and signs of these measures characterize the local convexity/concavity or the sphere-like *vs*. saddle-like character of the surface. Global shape, on the other hand, can be characterized by the so-called Minkowski functionals (MF). MF are versatile shape indices with strong roots in integral geometry, capable of robustly quantifying different aspects of spatial structure [2, 17, 18] (Figure 1). These shape indices, which can be of scalar or tensorial nature, are fundamental in the sense that they form a basis for any other additive functional that describes the shape of a 3D body (Hadwiger-Alesker theorems) [19–21].

**Figure 1:**
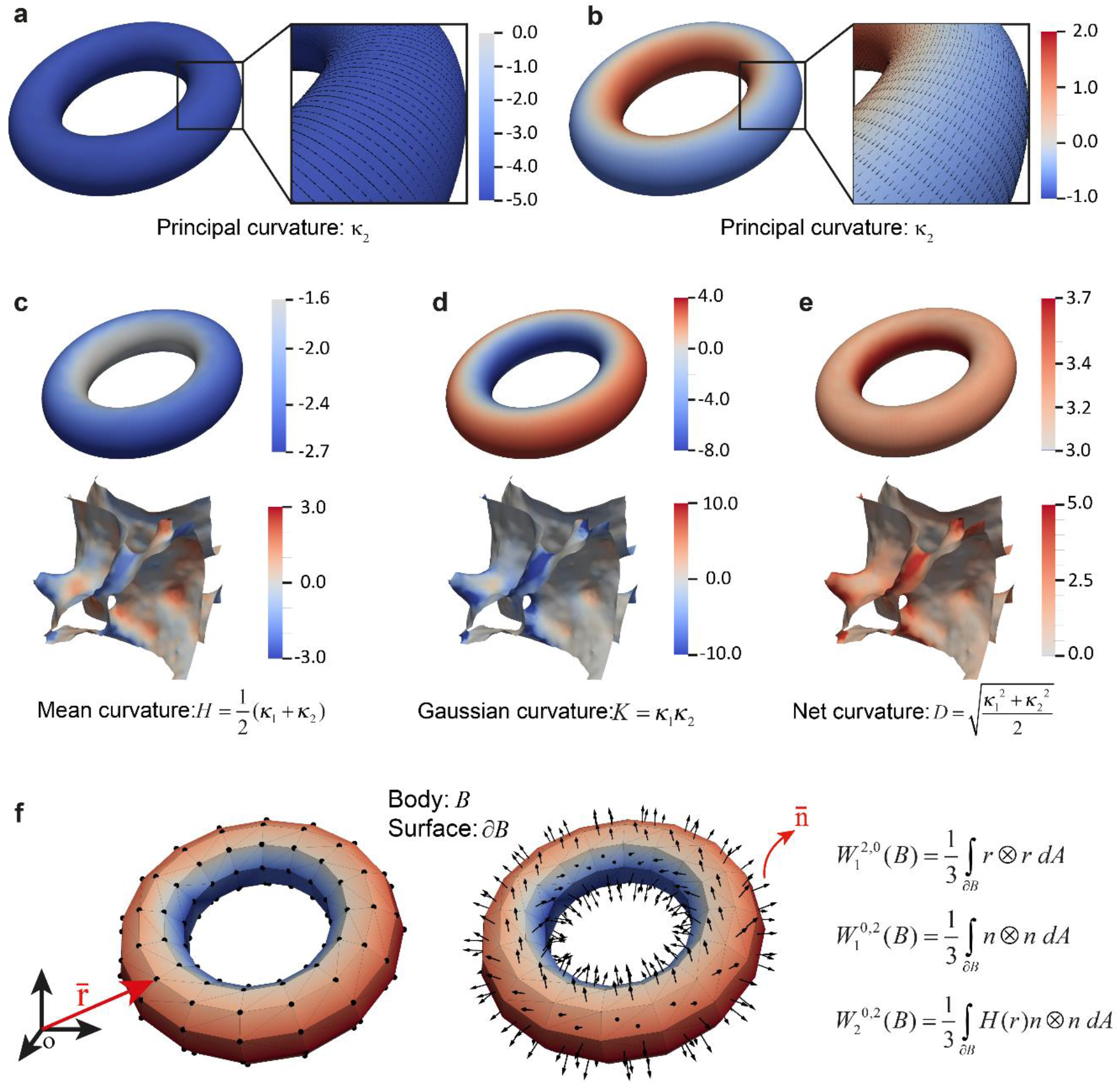
Surface curvature and Minkowski tensors. a-b) The minimum (*κ*_1_) and maximum (*κ*_2_) principal curvatures and associated principal directions on a torus model. c-e) The definitions of the mean (*H*), Gaussian (*K*), and net (*D*) curvatures as functions of the principal curvatures. The top row visualizes the curvatures of the torus, while the bottom row depicts some small sections of a trabecular bone interface. f) A visualization of the components used in the computation of the Minkowski tensors of a coarse torus model, showing the position vectors 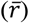 and normal vectors 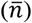, as well as the expressions for the tensors considered in this study.

Motivated by the desire for a unified description of morphology, we applied these local and global shape measures to hundreds of micro-computed tomography (micro-CT) scans obtained from bone biopsies of 70 donors at five anatomical sites [5]. At the local level, we computed the mean, Gaussian and net curvatures of the trabecular bone interfaces. We observed that the spatial curvature distributions are sensitive to differences in bone microarchitecture from different sites. At the global level, we computed the scalar and tensorial MF, and compared them with traditional morphometric indices. We focused on the more potent tensorial MF that can sensitively quantify the various types of intrinsic anisotropy, which is highly relevant in the study of trabecular bone [22, 23]. By using these fundamental shape descriptors, and reconciling them with previous metrics, we provide a novel geometric perspective on trabecular morphometry, that is compatible with virtually every other type of complex microstructure. Not only does this approach offer new insights into the trabecular morphology, we believe it could also complement the “language of shape” for the morphological design of synthetic tissue scaffolds [24].

## 2. Results

### 2.1. Surface curvature of the trabecular bone interface

We started our geometric analysis at the most local scale, by estimating the mean (*H*), Gaussian (*K*), and net (*D*) curvatures of the trabecular bone interfaces from their triangulated mesh representations. The mean curvature describes how much a surface is locally convex or concave. The Gaussian curvature quantifies the type of the surface: *K* < 0 signifies a saddle-shaped region (hyperbolic), *K* = 0 implies an intrinsically flat region (such as a plane, cylinder or cone), and *K* > 0 describes a sphere-shaped region (Figure 1). The net curvature is less common and describes how much a surface locally deviates from a planar region (Figure 1).

Figure 2 depicts the mesh representations of three representative trabecular bone specimens from the femoral head (FH), iliac crest (IC), and second lumbar vertebra (L2), color-coded by their curvature (representative visualizations of the calcaneus (CA) and fourth lumbar vertebra (L4) are provided in Supplementary Figure 1). The FH samples typically exhibited an apparently uniform dispersion of regions with positive and negative values of the mean curvature. Comparing this to the L2 specimen, we observed that the latter showed much more regions of highly negative mean curvature. This is the consequence of the many rod-like elements that are typically present in specimens from the lumber spine, as opposed to the primarily plate-like architecture in FH specimens [5]. The Gaussian curvature distributions clearly showed that the geometry of trabecular bone is, on average, hyperbolic in nature (*K* < 0). This has been reported before for a few bone biopsies [25]. This prevalence of negative Gaussian curvature is consistent with the high topological complexity (*i.e*., high genus) of trabecular bone, according to the Gauss-Bonnet theorem [26]. The net curvature captures regions where the trabecular surface is strongly bent, without distinguishing between the saddle- or sphere-like nature of these bends. In FH specimens, such regions corresponded primarily to arc-like transitions between plate-like elements, while high net curvature in IC or L2 specimens was concentrated in the cylindrically-shaped rod-like elements.

**Figure 2:**
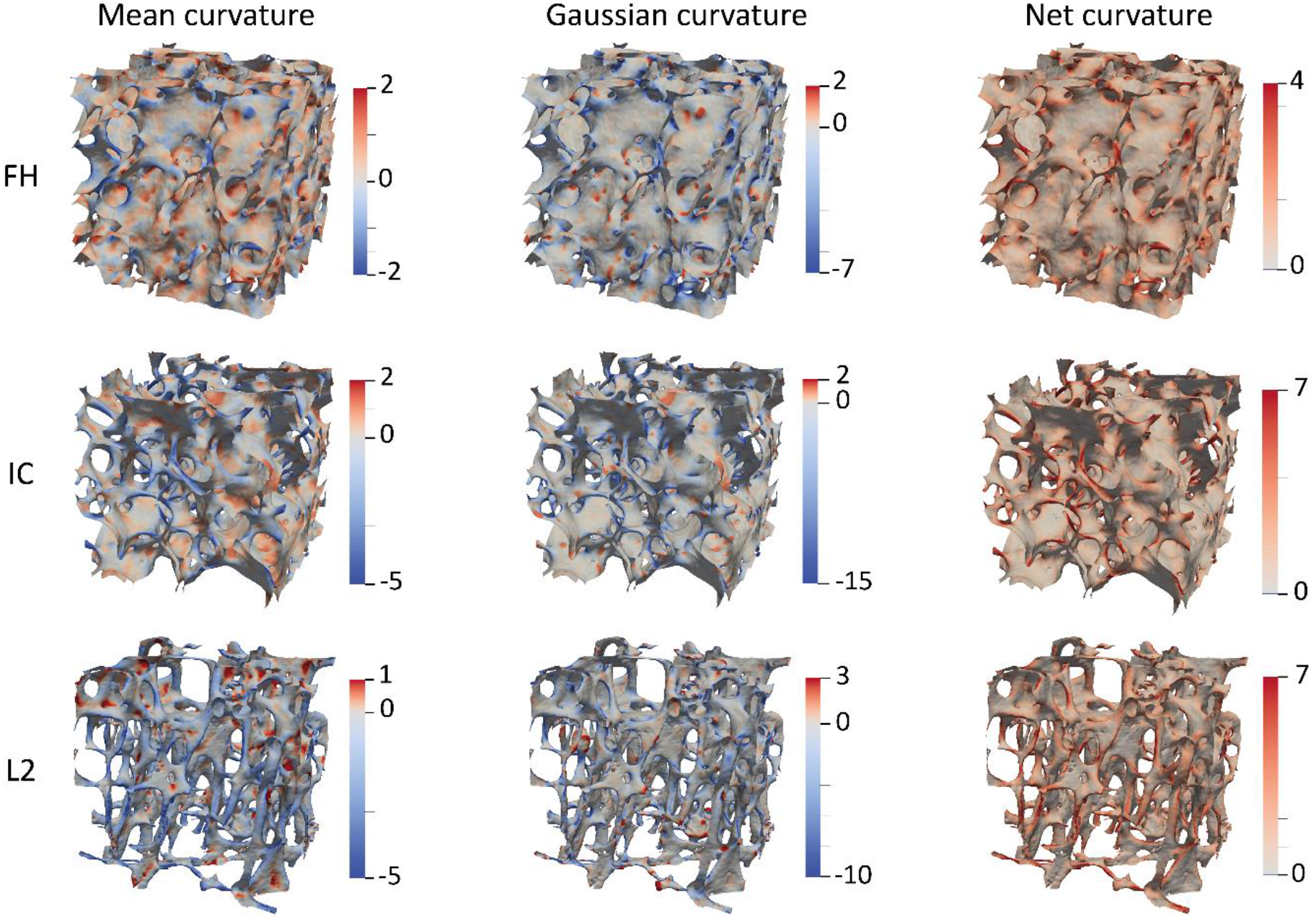
Curvature estimation on triangle meshes. The normalized mean, Gaussian, and net curvature visualizations of some representative (regarding morphology) trabecular samples from the femoral head (FH), iliac crest (IC), and second lumbar vertebra (L2).

### 2.2. Curvature distributions

Due to the inherently local nature of surface curvature, the average values of curvature are not of much descriptive use. In fact, the average mean and Gaussian curvatures are already captured in the structure model index (SMI~〈*H*〉) and Euler-Poincaré characteristic (χ~〈*K*〉), respectively [25]. Instead, it is important to consider the distribution of curvature throughout the trabecular specimens. Therefore, we computed the 1D and 2D probability distributions of the different curvature measures, obtained from more than 60 subjects at every anatomical location. The 1D probability densities of the mean curvature (Figure 3a) confirmed the above-mentioned observation that plate-like specimens (*i.e*., FH) exhibit a more uniform distribution of the normalized mean curvature, with a peak close to *H/S_v_* ≈ 0, than rod-like specimens (*i.e*., L2 and L4). The latter displayed a flattened and more negatively skewed distribution, centered around *H/S_v_* ≈ −0.5. The mean curvature density also reflected the intermediate nature of the iliac crest (IC) and calcaneus (CA) specimens, containing both plate-like and rod-like elements. As expected, the Gaussian curvature density functions were negatively skewed, with a sharp peak around 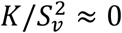. To assess the potential impact of age on curvature, we compared the curvature distributions of the specimens harvested from donors younger than 60 and those older than 80 for each anatomical site (Supplementary Figure 2). In the case of the mean curvature, significant differences in the probability distributions of both age groups were only observed for the L4 samples (two-sample Kolmogorov-Smirnov, *p* < 0.01). Indeed, the curve belonging to the donors older than 80 was more flattened and exhibited a lower peak value than the curve corresponding to donors younger than 60. These differences could be attributed to the progressive thinning and disappearance of thin rod-like elements, which is known to be more prevalent in the lumbar spine [27]. The Gaussian and net curvature profiles, however, did not show significant differences between age groups.

**Figure 3:**
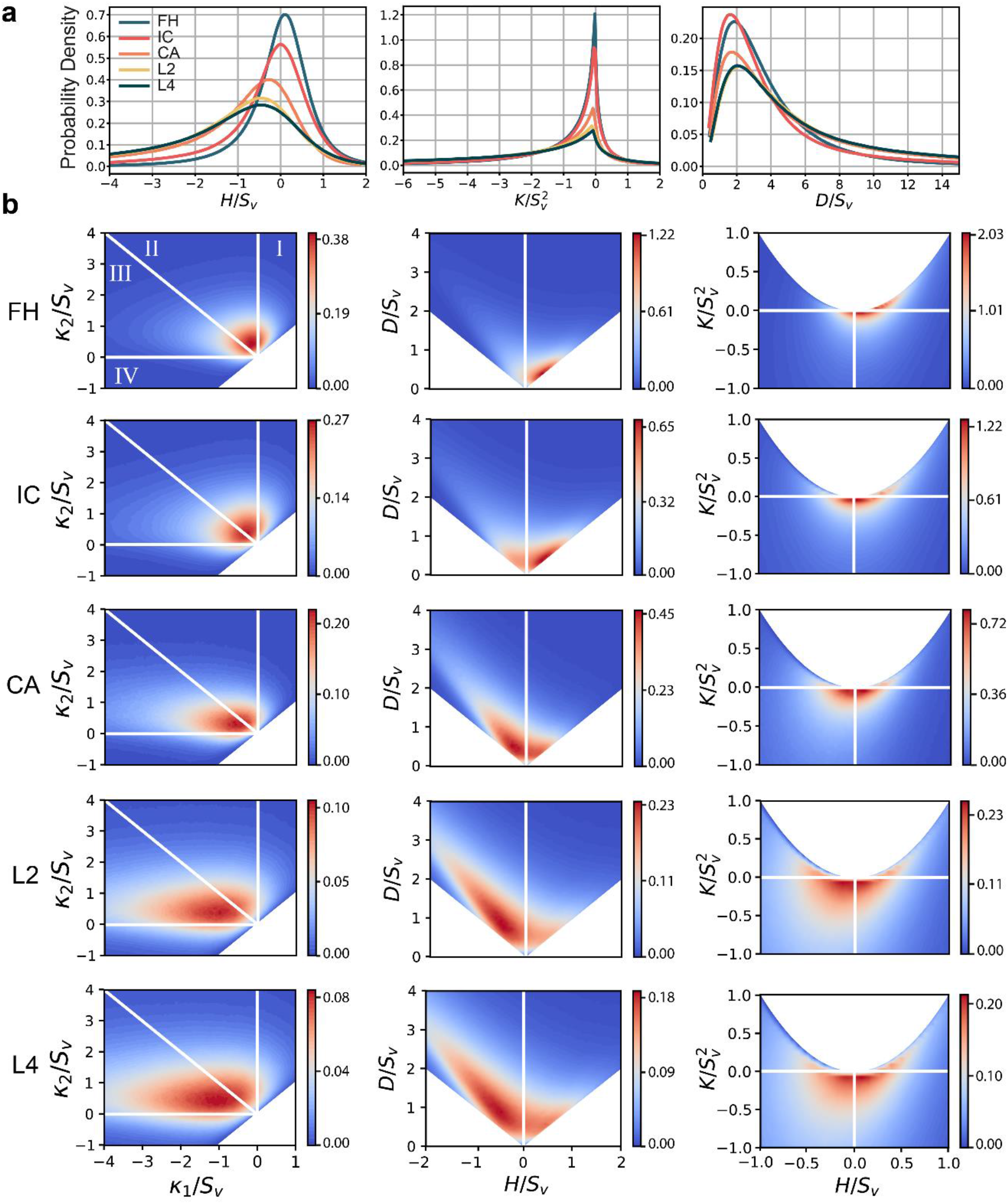
1D and 2D curvature distributions. a) Probability density distribution of the normalized mean (left), Gaussian (middle) and net (right) curvature per bone type. Each curve contains data from several samples (n_CA=66,n_FH=62,n_IC=68,n_L2=65,n_L4=68). b) The interface shape distributions (ISD) for specimens from every anatomical site, showing the normalized curvature probability densities. Left column displays the ISD of the principal curvatures (*κ*_1_ and *κ*_2_). Middle column displays the ISD of the net (*D*) and mean (*H*) curvatures. Right column displays the ISD of the Gaussian (*K*) and mean (*H*) curvatures. Each plot contains data from several specimens (*n_CA_* = 66, *n_FH_* = 62, *n_IC_* = 68, *n*_*L*2_ = 65, *n*_*L*4_ = 68).

Since a full description of surface curvature is typically built on two variables, such as the pair of principal curvatures, we quantified the interface shape distributions (ISD) for different curvature measures. These types of 2D probability density maps have been used to characterize the morphological evolution of spinodal decomposition systems during coarsening [28, 29]. The ISD of the principal curvatures is subdivided into different regions (Figure 3), providing an intuitive overview of the types of geometries that are encountered in the trabecular bone specimens. Saddle shapes appear in regions II and III, spherical shapes are situated in regions I and IV, and cylindrical shapes are found on the boundaries between the saddle-shaped and spherical regions (horizontal and vertical lines). For example, the principal curvature ISD of the FH specimens showed that most of the interface corresponds to saddle-shaped regions, and that sphere-like indentations (region I), and not protrusions (region IV), were the primary source of positive Gaussian curvature.

The ISD of the principal curvatures captured the progressive transition from primarily plate-like to primarily rod-like bone specimens (Figure 3). While the plate-like FH specimens exhibited a relatively concentrated, circular distribution of curvatures, the rod-like specimens of L2 and L4 were characterized by a much broader distribution with a horizontal orientation. This horizontal preference is due to the presence of rod-like elements in those specimens. These rod-like elements were not perfect cylinders, however, but were slightly saddle-shaped. The IC samples exhibited a principal curvature distribution that was similar to that of the FH, indicating primarily plate-like elements. For the CA samples, a horizontal orientation of the distribution was apparent, indicating a higher proportion of rod-like elements in the structures. The morphological differences between bone types were also observed in the joint probability distributions of the normalized net (*D/S_v_*) and mean curvature (*H/S_v_*), and the transition from plate-like to rod-like specimens was clearly visible (Figure 3). Moreover, these distributions again showed that the mean curvature of trabecular bone is, in general, not uniformly centered around zero. For our FH specimens, the peak of the mean curvature was situated slightly above *H* = 0, indicating shapes that were more concave than they were convex. This peak transitioned towards negative values for rod-like specimens, due to the convex nature of the rods. The ISD of the mean and Gaussian curvatures also captured this distinction between the different bone types, again showing a broader distribution for more rod-like specimens.

### 2.3. Radial distribution function

The ISD characterizes the local shape of trabecular bone by providing insight into the range and frequencies of the different types of curvature. However, it does not provide information about the way these curvatures are distributed in space and how the curvatures at different locations in the structure are related. Knowledge of this spatial correlation is relevant, since two structures could (theoretically) exhibit the same ISD, while having their curvatures distributed differently throughout space [30]. Therefore, we quantified the spatial correlation of the mean and Gaussian curvatures, using a curvature-based radial distribution function (RDF). Traditionally, the RDF has been employed in the analysis of granular systems, where it quantifies the likelihood of finding particles at a certain distance from a reference particle, relative to what would be expected based on the overall density of the system [31]. Depending on the type of the particle system and the associated interactions, the (excess) probability of finding neighboring particles will vary as a function of distance. The RDF has also gained popularity to quantify the spatial correlation in non-particulate systems [2]. When defined on the basis of the mean curvature, the RDF has been used to complement the ISD in characterizing the coarsening dynamics of spinodal decomposition systems [30, 32]. In this sense, the RDF provides a slightly more global interpretation of curvature than the ISD. Here, we used a similar approach to compute the RDF of the mean and Gaussian curvature for the trabecular bone specimens from the different anatomical sites. In case of the mean curvature, we define the RDF as:

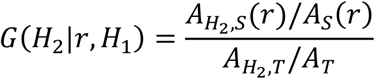

Here, *A*_*H*_2_,*S*_(*r*) is the total area of faces with a mean curvature of *H*_2_ within a spherical shell of radius r, centered around a reference point with a mean curvature of *H*_1_. *A_S_*(*r*) represents the total area of all faces inside the spherical shell, *A*_*H*_2_, *T*_ is the total area of faces with the mean curvature *H*_2_ in the entire specimen, and *A_T_* is the total face area of the entire specimen. Hence, *G*(*H*_2_|*r, H*_1_) describes the area-density of the faces with *H*_2_ at a distance r from a face with *H*_1_, relative to the overall area-density of the faces with *H*_2_. As such, the RDF captures how much more (*G* > 1) or less (*G* < 1) likely it is to find pairs of faces with a certain combination of mean curvature at a given distance from each other as compared to a random distribution throughout the specimen [32].

We plotted the RDF of the normalized mean (Figure 4) and Gaussian (Supplementary Figure 3) curvature at several characteristic distances. Taking the mean curvature RDF as the running example, the plots should be symmetric about the line *H*_1_/*S_v_* = *H*_2_/*S_v_*, since *G*(*H*_2_|*r,H*_1_) = *G*(*H*_1_|*r, H*_2_) [30]. Considering the RDF plots for 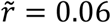 (left column in Figure 4), several observations could be made. For example, distinct positive correlations (*G* > 1) and anti-correlations (*G* < 1) were observed in all specimens. The positive correlations increased for the more extreme values of curvature, indicating that those strongly curved regions are highly concentrated in the structure. In other words, points with high mean curvature values are likely to have neighbors with high mean curvature as well. On the other hand, it is less likely to encounter neighbors with curvatures on the opposite sides of the spectrum (anti-correlation). For the FH, IC, and CA samples, a relatively stronger positive correlation was observed along the entire line *H*_1_ = *H*_2_ than for the L2 and L4 samples. Additionally, stronger anti-correlations were observed in the FH, IC and CA specimens. It is also noteworthy that the range of curvatures is substantially larger for the rod-like samples (L2 and L4) than for the plate-like samples (FH), indicating that the positive correlations in the rod-like samples occur at more extreme locations (relatively speaking) than for the plate-like samples. At the larger values of 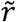 the correlations and anti-correlations gradually dissipated, and the RDF became more uniform and approached *G* = 1. The most extreme values of curvature maintained some positive correlation at 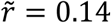, for all specimens. However, the correlation dissipation occurred faster in the L2 and L4 specimens, showing a more uniform distribution of *G* around unity than the FH, IC, and CA samples. The RDF of the Gaussian curvature (Supplementary Figure 3) exhibited a similar effect, although the positively correlated region had a more triangular shape. Moreover, at small and intermediate distances, high positive correlations were observed for the entire range of the positive Gaussian curvatures, indicating that locally spherical features are highly concentrated in trabecular bone.

**Figure 4:**
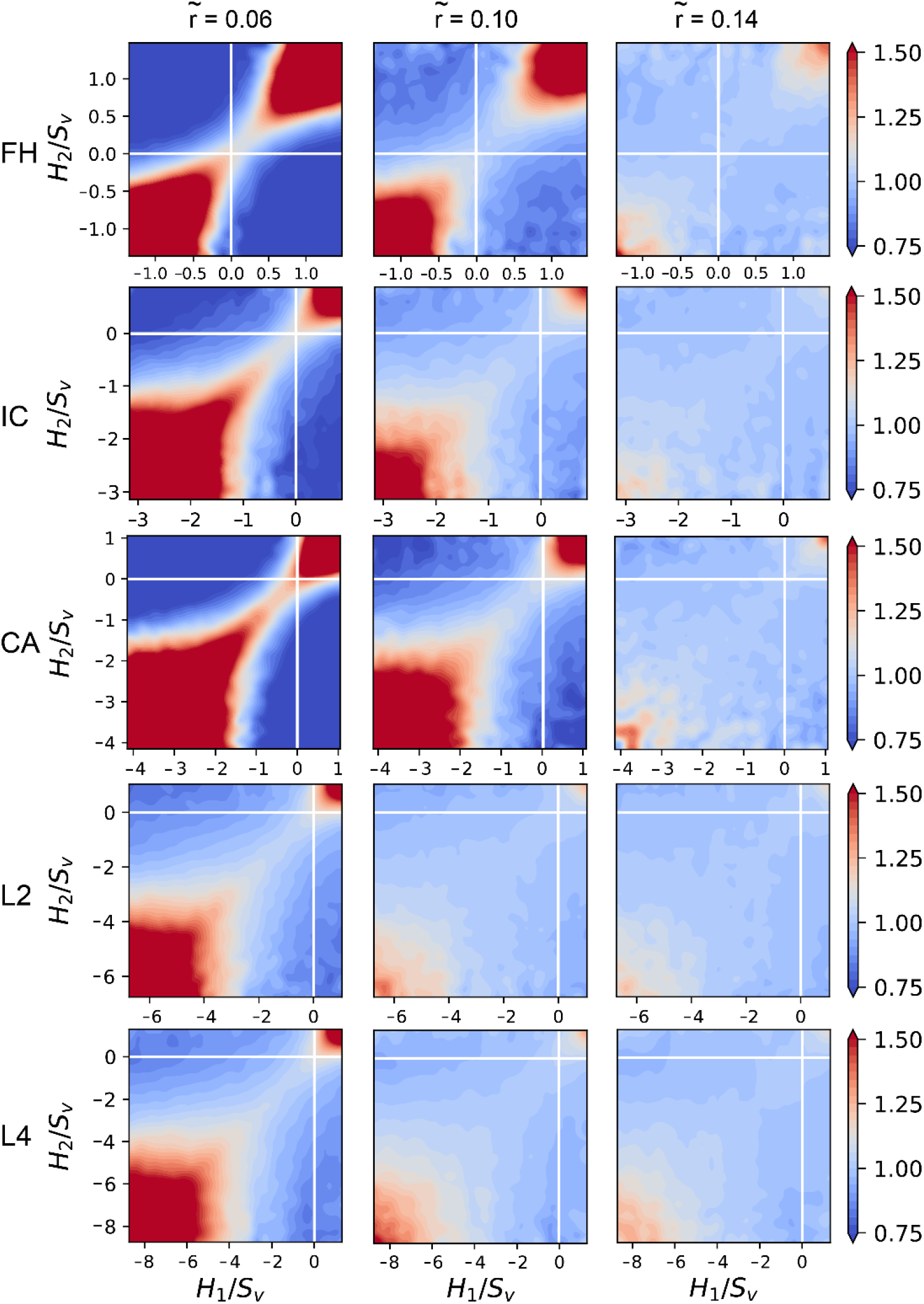
Radial distribution function. Plots showing the radial distribution function of normalized mean curvature (*G*) for representative (regarding morphology) specimens from the five anatomical sites, at different values of the characteristic distance 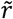. Color bar indicates enhanced (*G* > 1) or reduced (*G* < 1) probability of finding mean curvature pairs (*H*_1_ – *H*_2_) at a given distance 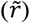 from each other, with respect to randomly distributed curvatures. Left column: 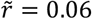, middle column: 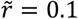, right column: 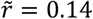.

### 2.4. Scalar Minkowski functionals

The ISD presents the most local measure of trabecular shape, by characterizing the curvatures at individual points along the interface. Two-point correlation functions, such as the RDF presented above, provide a slightly more global picture by considering pairs of points throughout the structure. Nevertheless, it is useful to complement these approaches with truly global, or integral, metrics that describe the shape as a whole. From an integral geometric viewpoint, the most fundamental indices to characterize the global shape are the Minkowski functionals. Their fundamental nature for shape description is described in Hadwiger’s theorem (generalized to tensors by Alesker), stating that any other motion-covariant, conditionally continuous, additive functional on a body is a linear combination of the Minkowski functionals [19–21]. In addition to their fundamental nature, Minkowski shape indices are also highly versatile, meaning that they can be applied to a broad spectrum of complex structures, and are robust against noise [2, 20].

The simplest types of Minkowski functionals are of a scalar nature and are further referred to as the Minkowski scalars (see Supplementary Note 1 for the mathematical formulations and background). For a 3D body *B* (Figure 1), four scalar MF can be defined, which are proportional to the volume (*W*_0_(*B*)), the total area of the bounding surface (*W*_1_(*B*)), the area-integrated mean curvature (*W*_2_(*B*)), and the area-integrated Gaussian curvature (*W*_(*B*)_). The latter is proportional to the Euler-Poincaré index, a topological invariant describing connectivity. The Minkowski scalars have been applied in the analysis of various spatial architectures, including voxelized representations of trabecular bone [33]. Here, we computed the Minkowski scalars *W*_1_, *W*_2_, and *W*_3_ on the smoothed triangle meshes of the trabecular bone interface, and compared them to traditional bone morphometric indices that characterize the global trabecular shape (Figure 5) [5, 12]. The scalar *W*_0_ was omitted, since it is not defined for open surfaces. The scalar *W*_1_ and the bone surface area (BS) were relatively well correlated (Figure 5a). The strongest correlation was observed for the L4 specimens (ρ = 0.91), while the weakest correlation was attained in the CA specimens (*ρ* = 0.66). The deviations between the computed *W*_1_ and BS could potentially be attributed to different underlying meshes (BS was directly calculated on a marching cubes mesh [5]). The scalar *W*_2_, which captures the area-integrated mean curvature, is plotted against the morphometric parameter *∂S/∂r*, showing strong correlations for all bone types (*ρ* > 0.87, Figure 5b). The parameter *∂S/∂r* represents a surface area derivative, and is estimated as the change in surface area (*dS*) when the surface is dilated by a small amount, divided by the length of that dilation (*dr*). The area of such a dilated parallel surface (*S_r_*) is related to the area of the original surface (*S*_0_) and its curvature by [25]:

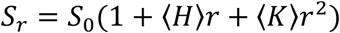

where *r* is the signed distance from the original surface, 〈*H*〉 is the average mean curvature, and 〈*K*〉 is the average Gaussian curvature. The dilation-based parameter *∂S/∂r* appears in two well-known bone morphometric indices: the SMI and the 3D trabecular bone pattern factor (TBPf) [13, 34, 35]:

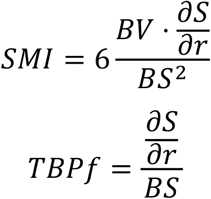

**Figure 5:**
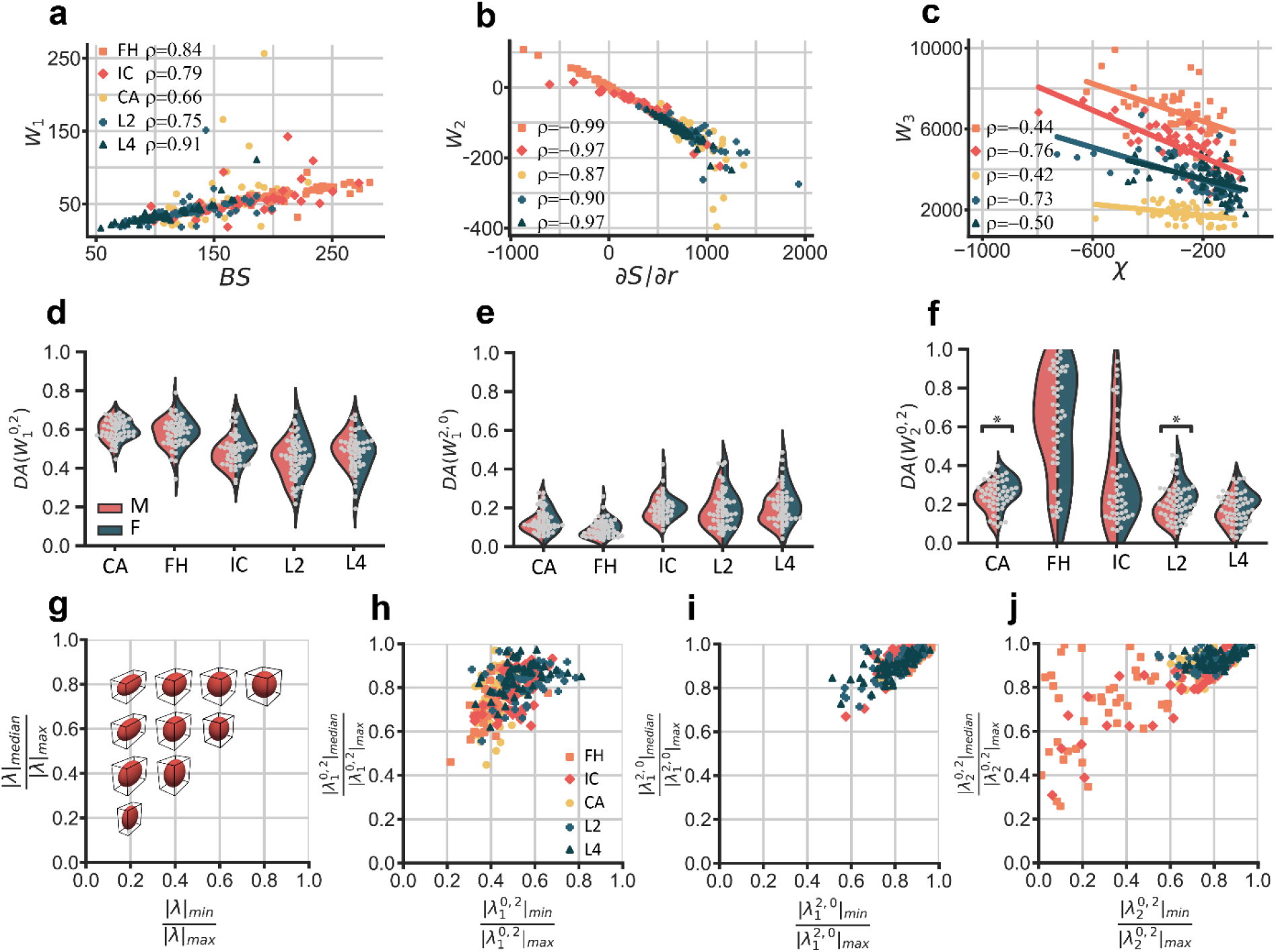
Scalar and tensorial Minkowski functionals applied to trabecular bone. a-c) Results for the three Minkowski scalars, plotted versus their equivalent standard morphometric index: a) *W*_1_ versus bone surface (BS), b) *W*_2_ versus surface area derivative (*∂S/∂r*), c) *W*_3_ versus Euler-Poincaré characteristic (*χ*). d-f) The distribution plots for the degree of anisotropy (DA) with respect to the different Minkowski tensors, plotted per anatomical site and split between the specimens from male (M) and female (F) donors: d) DA for tensor 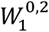, e) DA for tensor 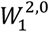, and f) DA for tensor 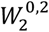. *:*p<0.01*. g-j) Ellipticity with respect to the different Minkowski tensors, shown as the ratio of the median to the maximum eigenvalue versus the ratio of the minimum to the maximum eigenvalue: g) an illustration of the various degrees of ellipticity, h) ellipticity with respect to 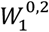. i) ellipticity with respect to 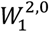, and j) ellipticity with respect to 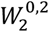.

In that sense, both SMI and TBPf are proportional to the average mean curvature (〈*H*〉) of the surface (for small dilations (r^2^≈0), the second-order Gaussian curvature contribution can be neglected), essentially meaning that the 〈*H*〉 is a global morphometric index [25, 27]. Overall, the correlation between *W*_3_ and the Euler-Poincaré index (*χ*) is lower (0.42 < *ρ* < 0.76) than the correlations between the previous Minkowski scalars and their corresponding morphometric indices (Figure 5c). This could again be attributed to the different calculation approaches: *W*_3_ is based on the integral Gaussian curvature of the triangle meshes, while *χ* is computed on 3D binary images. Finally, an important observation is that *W*_3_ captures the differences between the specimens from different anatomical sites, while these are not reflected in the *χ* values (Figure 5c). This implies that *W*_3_ could potentially be more sensitive to subtle changes in connectivity than *χ, e.g*. in case of disease.

### 2.5. Tensorial Minkowski functionals

A relatively novel extension of the Minkowski scalars for global shape quantification is provided by the so-called Minkowski tensors (MT). Due to their tensorial nature, these MT capture the orientation-dependent aspects of morphology, a feature that is highly relevant for the study of heterogeneous materials such as trabecular bone [22, 23]. While the MT have been employed to characterize granular packings, galaxies, or foams [17, 36, 37], we are the first to apply these tensors to the study of trabecular bone. As a natural consequence of their mathematical foundation, many different MT can be defined, each characterizing a different aspect of morphology. In principle, the MT can be defined for any arbitrary rank, but we primarily focused on rank-two tensors, due to their intuitive physical interpretation [2]. Higher rank MT are briefly considered in section 2.6. For a 3D body, six relevant rank-two MT are defined (Figure 1 and Supplementary Note 1). As an example, the tensor 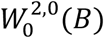 is a measure of the spatial distribution of mass for a solid body *B*, in some sense analogous to the moment of inertia tensor. The tensor 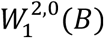, on the other hand, measures the mass distribution when the entire mass of *B* is homogeneously distributed on the surface (*i.e*., a “hollow” body). Here, we considered the aforementioned (translation-covariant) tensor 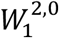 as well as two other (translation-invariant) Minkowski tensors, namely 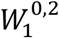 and 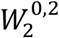. The tensor 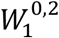 describes the distribution of the surface normal vectors, while 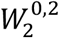 describes the distribution of the mean curvature (surface normals weighted by curvature).

Every MT can be used to quantify anisotropy with respect to that particular tensor. We define the degree of anisotropy (DA) for a tensor 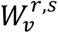 as:

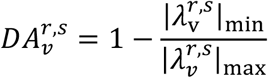

Here, 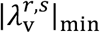 and 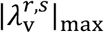 are the absolute values of the minimum and maximum eigenvalues of the tensor 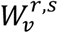. As such, we were able to quantify the different types of anisotropy of the trabecular bone samples, including the anisotropy of the interface orientation 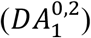 or the anisotropy of the mean curvature 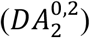. For trabecular bone, a classical and popular approach to quantify anisotropy has been based on the mean intercept length (MIL) method [12, 38, 39]. However, the MIL is limited to interfacial anisotropy (by definition) and is known to suffer from some conceptual shortcomings, such as noise sensitivity, sampling bias, and poor data fitting in specific cases, that could invalidate the anisotropy results [40]. Instead, Minkowski tensors are robust alternatives, showing higher sensitivity to anisotropy in 2D boolean model systems [41]. To assess the potential of Minkowski tensors as alternatives to the MIL tensor for bone anisotropy quantification, we computed the DA of both approaches on 259 trabecular bone specimens. The relevant Minkowski tensor for this comparison is 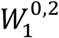, as it describes the interfacial orientation. We found that, with the most recent algorithm for the calculation of *DA_MIL_* (in BoneJ [42]), all the data for *DA_MIL_* and 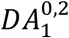 were strongly correlated (Spearman’s *ρ* = 0.957, Pearson’s *ρ* = 0.963) and centered around the identity line (Supplementary Figure 4). Both methods also predicted similar principal directions with most angle differences below 10° (Supplementary Figure 4b). In contrast to theoretical predictions on 2D model systems, our analysis indicates that both approaches yield similar results on high-resolution trabecular bone scans [41]. Despite these similar results, it must be emphasized that *DA_MIL_* is highly dependent on the specific MIL implementation, which has been shown to be a potential source of significant variation in classical MIL algorithms [15, 43, 44]. Moreover, the MT approach is inherently less sensitive to noise and more computationally efficient than the MIL approach, since it does not rely on counting lines and intersections [41], and also offers the ability to quantify the other types of anisotropy.

Comparing the 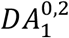 distributions (Figure 5d), we observed that all bone types exhibit a distinct level of interfacial anisotropy, with significant differences between the means of the different bone types (Supplementary Table 1). Higher mean values were obtained in the 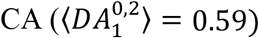 and 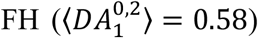 specimens, as opposed to the 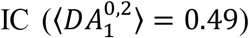, the 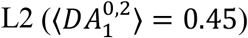, and 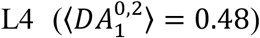 samples. Moreover, a wider spread in the anisotropy values was observed in the L2 and L4 specimens. In all cases, the degree of anisotropy with respect to the tensor 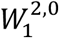, which characterizes the mass distribution of the “hollow” trabecular bone, was much lower (Figure 5e). Significant differences between the different bone types were detected, and higher mean values were attained for the 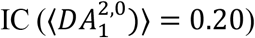, the 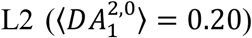 and the 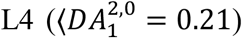 samples as opposed to the 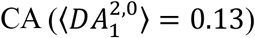 and the 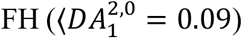 samples. Finally, 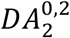 quantifies the anisotropy of the curvature-weighted surface normals (Figure 5f), again displaying significant differences in the means between the bone types. The 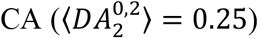, 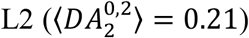, and 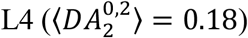 specimens exhibited narrower distributions than the 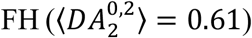 and 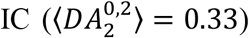 specimens. Interestingly, there were statistically significant differences between the mean 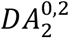 values calculated for the specimens harvested from male and female donors in the case of the CA (*n*_1_ = 25, *n*_2_ = 27, Mann-Withney U = 179, *p* = 0.004) and L2 (*n*_1_ = 25, *n*_2_ = 30, Mann-Whitney U = 179, *p* = 0.001) bone types.

In reporting DA, only the extremal tensor eigenvalues were considered. To extend our characterization of the Minkowski tensors, we also plotted the ratio of the median to the maximum eigenvalue against the ratio of the minimum to the maximum eigenvalue. Since rank-2 tensors can be represented by the surface of an ellipsoid (Materials & Methods), these plots provide insight into the “ellipticity” of the bone specimens with respect to a particular tensor (Figure 5g). Data for which 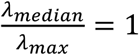 are represented by prolate spheroids, while data on 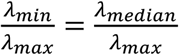 are represented by oblate spheroids. When 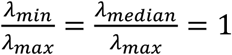, a perfect sphere is obtained and the data is considered fully isotropic with respect to that particular tensor. For the tensor 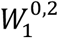, the data was clustered between the oblate and prolate shapes on the ellipsoid spectrum (Figure 5h). Moreover, the ellipticity of the 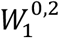 data was in good agreement with the ellipticity of the MIL data (Supplementary Figure 4d). In the case of the 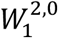 tensor, most of the specimens were highly concentrated in the nearly isotropic region, with some data points (IC, L2, and L4) exhibiting higher ellipticity (Figure 5i). For the 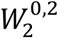 tensor, the data for the CA, L2 and L4 specimens was again concentrated in the nearly isotropic region, but the data for the FH and IC specimens were scattered over the entire ellipsoid spectrum. For example, the FH specimens covered both highly oblate and prolate ellipticity, with various degrees of anisotropy (Figure 5j). Taken together, these plots underscore that interfacial orientation is only one of several sources of bone anisotropy and ellipticity, and that other sources can be quantified by considering a different Minkowski tensor (e.g. 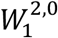 or 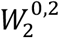).

### 2.6. Anisotropy in spatially decomposed bone

The Minkowski functionals provide a global (integral) interpretation of trabecular shape, by assigning either a scalar or a tensor to the entire region of interest. However, the shape and size of this region could be chosen arbitrarily within the cubic specimen volume. Hence, it is possible to apply the Minkowski analysis to several smaller substructures, in order to create a Minkowski map that quantifies the intra-specimen variations of the integral shape indices [2]. In that sense, such a spatially decomposed analysis of the Minkowski functionals occupies an intermediate position between the highly localized analysis of curvature distributions and the whole-specimen shape characterization presented in Sections 2.3 and 2.4.

To quantify the intra-specimen anisotropy changes, we decomposed 100 trabecular specimens into a set of smaller components. In order to maintain representative trabecular substructures, we used a 3×3×3 cubic grid for this spatial subdivision (Supplementary Figure 10b). We computed the two translation-invariant Minkowski tensors 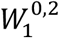 and 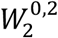 on the resulting 2700 substructures, enabling a local characterization of the ellipticity with respect to those tensors. We found that, in general, the ellipticity varies throughout the specimens and is different for both tensors (Figure 6a-d). To quantify the spatial variation, we calculated the relative difference between the anisotropy of a substructure and that of the entire specimens (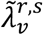, Materials & Methods), as well as the angle difference between the local and global principal orientations 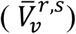. For both tensors, the local DA varied substantially with respect to the whole-sample value and led to different distributions for both tensors (Figure 6e). Distinct angle differences in the local and global principal directions were also observed for both tensors, and wider variations were detected in the L2 and L4 specimens (Figure 6f).

**Figure 6:**
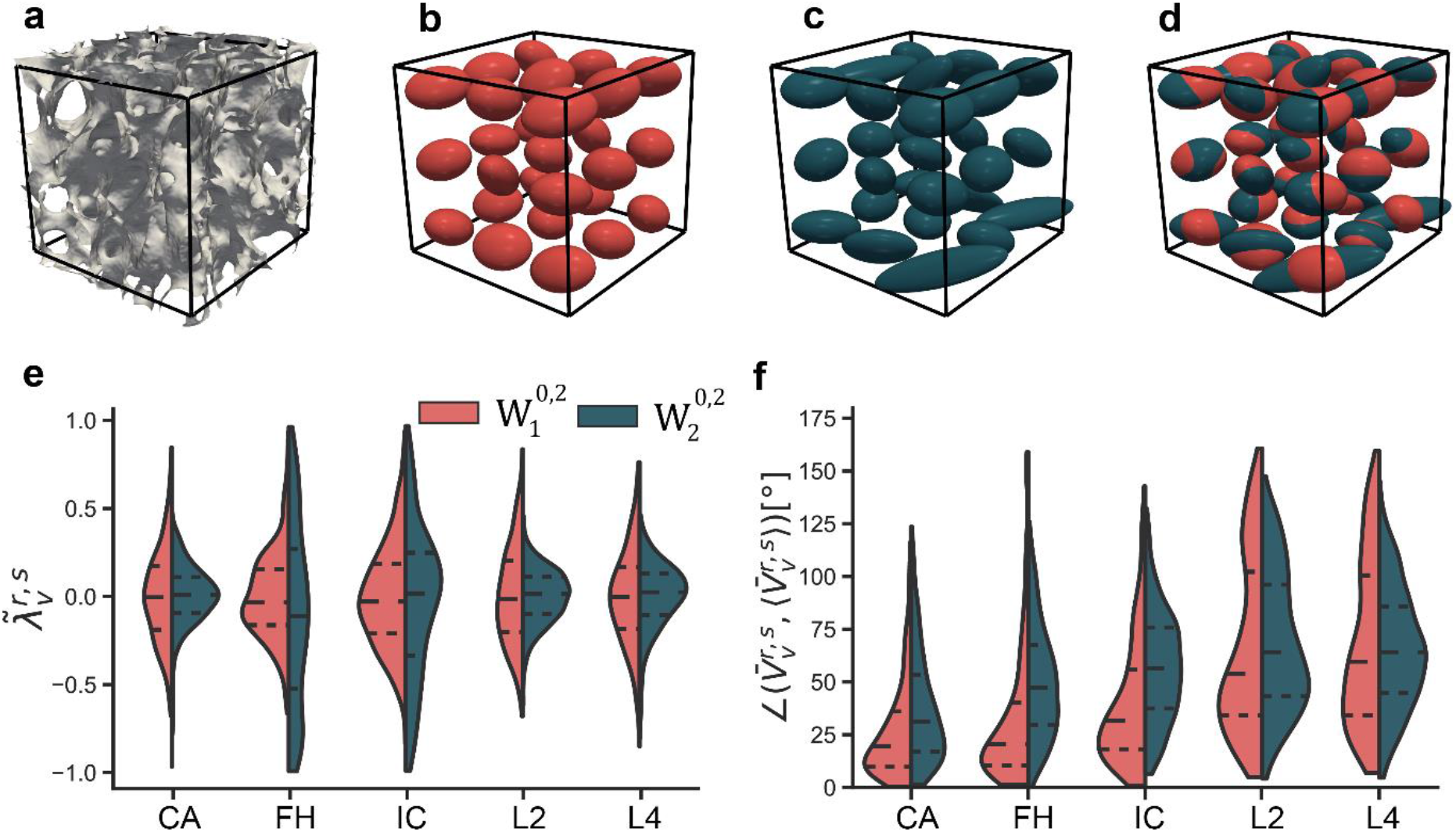
Minkowski tensor analysis on spatially decomposed samples. a) A visualization of a full FH trabecular bone specimen. b) Ellipticity with respect to the normal density tensor 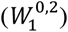 of the sample in a) in 27 subdomains. The ellipsoids are oriented in their principal direction. c) Ellipticity with respect to the curvature density tensor 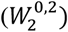 of the sample in a) at 27 subdomains. d) An overlay of the ellipticities of b) and c). e) The relative differences in the local and global anisotropy for 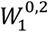 and 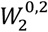, plotted for each anatomical site. f) The differences in the local and global principal orientations for 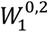 and 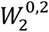, plotted for each anatomical site.

Finally, we asked whether higher-rank Minkowski tensors (beyond rank two) could provide additional insight into the structural and anisotropy differences between the different bone types. To this end, we calculated the quadratic (*q_s_*) and cubic (*w_s_*) rotational invariants of the so-called irreducible Minkowski tensors (Supplementary Note 1) for the spatially decomposed specimens. This analysis was motivated by the recent results in particulate matter, where these scalar invariants have been used as efficient structure metrics to detect local crystalline states in disordered packings of convex shapes [45, 46]. Due to their higher-rank nature, however, the physical significance of these structure metrics is less easily understood than for the rank-2 Minkowski tensors. Plotting the probability distributions of *q_s_* and *w_s_* for the FH and L4 specimens (540 data points each, Supplementary Figures 5 and 6), we observed globally smooth distributions for *q_s_* and *w_s_* that are qualitatively similar to those obtained for hyperuniform amorphous cellular solids [46]. Sharp peaks in the distribution would indicate the presence of a locally crystalline region with a certain structural symmetry. Significant differences between the structure metric distributions of the FH and L4 specimens were detected (two-sample Kolmogorov-Smirnov, *p* < 0.01), except for *q*_5_, indicating that these higher-order structure metrics are sensitive to the structural differences between the plate-like and rod-like specimens.

## 3. Discussion

The aim of this study was to provide a more fundamental geometric viewpoint on the quantification of trabecular bone shape. At the local perspective, this could be accomplished by quantifying the surface curvature of the trabecular interface. In fact, curvature is the defining characteristic when it comes to distinguishing between local structural features, such as rods (*K* = 0, *H* < 0), plates (*K* = 0, *H* = 0), and saddle-shaped arcs (*K* < 0), or to identify primarily convex (*H* < 0) or concave (*H* > 0) regions. We quantified the complex curved landscapes of trabecular bone using the ISD, finding that these density maps serve as effective shape fingerprints for trabecular bone from different anatomical sites. Indeed, the ISD captured the morphological differences between plate-like (FH) and rod-like (L2 and L4) specimens, but were also sensitive to intermediate morphologies along the plate-rod spectrum (IC and CA). Interestingly, the ISD also revealed that trabecular bone is not approximating a minimal surface, opposing a claim often made in the literature to support the use of minimal surface-based scaffolds. Minimal surfaces, for which *H* = 0 everywhere, arise in systems where surface energy is minimized (*e.g*., soap films). It is often assumed that a similar phenomenon takes place in the formation and remodeling of trabecular bone, leading to an overall minimal surface morphology. However, we find that the principal curvature ISD of trabecular bone is distinctly different from the minimal surface ISD, where all data points lie on the boundary between regions II and III (*κ*_1_ = −*κ*_2_, Figure 3b).

We also performed a global shape analysis of the trabecular bone interface. To this end, we employed the scalar and tensorial Minkowski functionals, since these are fundamental, highly versatile, and robust indices for integral shape quantification. We found that the Minkowski scalars, which were computed directly on the triangulated bone meshes, correlated with traditional bone morphometric indices, such as BS, *∂S/∂r*, or *χ*. Moreover, we found that *W*_3_ was more sensitive to differences in bone microarchitecture than the corresponding traditional metric *χ*. Our work was the first to apply the Minkowski tensors to the quantification of trabecular bone shape. This analysis revealed different degrees of anisotropy and ellipticity, depending on the morphological aspect that is being considered. Moreover, anisotropy differences between bone specimens harvested from different anatomical sites could be detected. An important aspect of this Minkowski functional approach is that it unifies several traditional morphometric indices within the same geometrical theory. For example, interfacial and volume anisotropy are traditionally characterized using different methods (*e.g*., the MIL and SVD methods), while both can be described within the Minkowski tensor framework by using a different tensor. We also applied higher-rank Minkowski metrics to the shape quantification of spatially-decomposed bone specimens, showing that they are also sensitive to morphological differences in bone from different anatomical sites. However, we note that these higher-rank metrics are usually applied to disordered assemblies of discrete convex bodies, such as the Voronoi diagram of a granular packing [45, 46]. Such materials are naturally well-suited for this type of domain-wise analysis, since the basic definition of these structure metrics is centered around decomposing the normal density of convex bodies into spherical harmonics. As such, analyses using these metrics might be more compatible with convex particle systems than with non-convex, smooth trabecular bone structures. In this regard, it would be interesting to apply this analysis to specimens that are decomposed into (almost) convex units, for example by volumetric decomposition into rods and plates [34], or by approximate convex decomposition [47]. Additionally, the use of clustering algorithms (*e.g*. k-means or DBSCAN) could shed light on the classification sensitivity of such higher-order metrics.

The key characteristic of our metrics is their fundamental geometric nature, which offers a unifying view and geometrical foundation for traditional bone morphometric indices. This geometric perspective could advance the understanding of morphological changes in aging and disease, such as the elusive plate-to-rod transition in osteoporosis [48]. Our approaches could also provide a framework for shape description within the context of bone healing, for example, to characterize the structure of the developing callus *in vivo* [49, 50]. Since these metrics are not bound by scale, they could also readily be applied to high-resolution images of much smaller structures within bone, such as the lacuno-canalicular network [51]. Moreover, our insights into trabecular bone curvature are relevant for recent investigations into the role of substrate curvature as a mechanobiological cue at the cell and tissue levels [7, 52], which could be leveraged in tissue engineering applications. In summary, we have provided a geometric approach to trabecular bone morphometry, quantifying both the local and global shape of trabecular bone, and unifying several traditional morphometric indices within the mathematical language of geometry. Our analyses were centered around surface curvature and Minkowski functionals, which proved to be sensitive fingerprints to site-specific differences in bone morphology. These approaches could facilitate the geometrical characterization of a broad spectrum of spatially-complex materials beyond bone, ultimately advancing the development of accurate structure-property relationships for such materials.

## Acknowledgments

The first author is thankful to Fabian Schaller and Sebastian Kapfer for sharing insights on the use and interpretation of the Karambola software, which was used for the computation of the Minkowski functionals, to Prof. Amber Genau and Delphine De Tavernier for useful discussions on the implementation of the radial distribution functions, and to the IDEA League program for facilitating an exchange at ETH Zurich. The research leading to these results has received funding from the European Research Council under the ERC grant agreements no. [677575] and [741883].

## 4. Materials & Methods

### 4.1. Trabecular bone data set

All analyses were performed using previously-published, high-resolution μCT data from the European Union BIOMED I Concerted Action “Assessment of Bone Quality in Osteoporosis”. The details of the database composition and data acquisition protocols can be found elsewhere [5, 53]. Briefly, the data set comprised trabecular bone biopsies harvested from 70 donors (38 male, 32 female, age between 23 and 92 years) at five anatomical sites: the femoral head (FH), the iliac crest (IC), the calcaneus (CA), the second lumbar vertebra (L2), and the fourth lumbar vertebra (L4). The specimens were scanned using a high-resolution μCT system (μCT 20, Scanco Medical AG, Switzerland) with a spatial resolution of 28 *μm* and cubic voxels with 14 *μm* length (for the CA samples, cubic voxels with an edge length of 28 *μm* were used). A 4 mm^3^ cubic volume of interest (VOI) was selected from the resulting scanned data, and 3D binary images of the mineralized bone phase were obtained after Gaussian filtering and thresholding. These binary images served as the basis for all consecutive analyses.

### 4.2. Surface reconstruction

The binary volume data was processed using custom Python codes and Python-based mesh processing libraries [54, 55]. First, a triangle mesh of the trabecular bone surface was reconstructed from the volume data using a marching cubes algorithm with a spacing equal to the voxel size (Supplementary Figure 7a) [56]. No padding was applied to the volume data prior to the marching cubes algorithm, ensuring that only the trabecular interface was meshed and resulting in a mesh that was a 2-manifold with boundary (Supplementary Figure 7c). Next, degenerate (zero-area) faces and small disconnected components were removed from the mesh. To account for the roughness inherent in marching cubes meshes, the triangle meshes were smoothed using implicit fairing. This is an efficient smoothing approach that is based on the implicit integration of a diffusion process, and guarantees volume preservation during smoothing [57]. The algorithm solves the linear system:

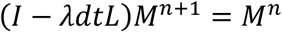

where *M^n^* represents the set of mesh vertices at iteration *n, L* represents the Laplacian, and *λdt* is a user-defined smoothing constant. The trabecular bone meshes were all smoothed using a single iteration and *λdt* = 5, enabling the smoothing of the marching cubes artefacts while maintaining small curved features on the trabecular bone surfaces.

### 4.3. Curvature estimation algorithm

The principal curvatures of the trabecular bone surface were estimated at every vertex of the discrete triangle mesh using an efficient multiscale fitting algorithm [54, 58]. The applied algorithm was an adaption of the Osculating Jets method [59], and fitted a second-order polynomial to a local neighborhood around every vertex for the curvature estimation. The local neighborhood was defined as a ball with a user-defined radius, which was centered at the vertex of interest. For the FH, IC, L2, and L4 trabecular bone meshes, the radius was defined as:

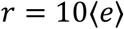

where 〈*e*〉 represents the average mesh edge length for a particular bone specimen (〈*e*〉 ≈ 20 μm). In the case of the CA meshes, which were scanned with a larger voxel size, the radius was set to *r* = 5〈*e*〉, corresponding to a similarly-sized neighborhood as compared to the other bone specimens. The sensitivity of the curvature estimation algorithm to the neighborhood size (*r*) and smoothing parameter (*λdt*) was assessed by calculating the interface shape distributions for a representative sample at different combinations of *r* and *λdt*, and by visualizing color-coded curvature distributions on the triangle meshes (Supplementary Figures 8 and 9).

### 4.4. Curvature probability density distributions

All reported curvatures were non-dimensionalized using a characteristic length parameter [32]:

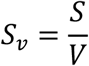

where *S* is the total mesh area and *V* = 4^3^*mm*^3^ is the volume of the cubic specimen. Face curvature values were calculated by averaging the curvatures of the three vertices associated with every face. In constructing the probability density distributions of curvature, the face curvatures were weighted by their face area. In case of the interface shape distributions (ISD), this implied that the ratio of the face areas with a certain combination of curvature to the total mesh area was considered. For example, in case of the ISD of the principal curvature, this means [28]:

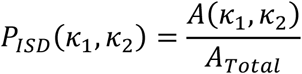

### 4.5. Radial distribution function of curvature

To compute the radial distribution functions (RDF) of the mean and Gaussian curvatures, the curvature values in the range of the 0.5 percentile and the 99.5 percentile were considered. These curvature values were binned in 100 bins of equal width and were weighted by their face areas. Since the RDF considers the curvatures of face pairs separated by a given distance, a characteristic distance was defined in function of the previously described characteristic length scale *S_v_*:

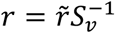

where 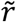 is a user-defined length parameter. In order to find face pairs, a spherical shell of nominal radius *r* and thickness Δ*r* was centered around the barycenter of every sample face of interest, where:

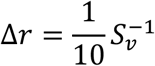

All faces with barycenters inside the spherical shell were considered as paired faces to the sampled face. For every bin, 1000 unique random faces were selected and the curvatures of the corresponding paired faces were computed and stored in area-weighted histograms with 100 bins of equal width. If the bin contained less than 1000 faces, which could occur at the extreme values of the curvature range, all available faces were used as sample points in the RDF. By sampling every bin, and combining the corresponding area-weighted curvature histograms of the paired faces, the RDF was constructed as a 100 × 100 matrix with probability values.

### 4.6. Minkowski structure metrics

The Minkowski functionals (scalars *W_v_*, tensors 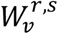) and the rotational invariants of the irreducible Minkowski tensors (IMT) (*q_s_* and *w_s_*) were computed on the trabecular triangle meshes using a C++ code (https://github.com/morphometry/karambola) that iterates over all faces, edges, or vertices to compute the relevant Minkowski metrics, in accordance with Table 2 of reference [20]. In order to prepare the trabecular bone meshes for the Minkowski metric computation, the non-manifold edges and vertices that could result after the marching cubes reconstruction had to be removed. The non-manifold edges were removed by constructing the face adjacency matrix of the mesh and removing those faces with edges shared by more than two faces. Non-manifold vertices that remained after the deletion of non-manifold edges were removed using Meshlab [60]. Of the six relevant Minkowski tensors, only the tensors 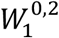 and 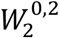 are translation-invariant tensors, which means that:

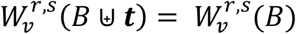

where 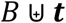 signifies the translation of body B along vector ***t*** [20]. The other Minkowski tensors are translation-covariant, and do not satisfy this relationship. For those translation-covariant tensors, a consistent definition of the mesh origin is required to enable a fair comparison between the different meshes. For all the trabecular bone meshes, the origin was defined to be in the center of the cubic bounding box.

In order to deal with the open trabecular bone meshes, a domain-wise analysis of the Minkowski metrics was performed. To this end, the faces at the boundaries of the mesh were assigned a different label than the faces inside the trabecular surface, and the Minkowski metrics were computed for each labeled domain separately. In this way, the faces not on the boundary are effectively considered to be part of a “closed” surface, and all the relevant Minkowski metrics could be calculated, which is not the case for the “open” boundary faces [20]. Representing a mesh as a doubly-connected edge list (DCEL), the boundary faces are those faces with at least one half-edge that appears only once in list (*i.e*., it is not shared with another face). A visual representation of the boundary face labeling is provided in Supplementary Figure 10a.

### 4.7. Spatial decomposition

A domain-wise Minkowski analysis was performed on spatially decomposed meshes. The mesh was subdivided into a 3×3×3 cubic grid, and all faces were assigned a label based on the location in the grid, resulting in a total of 28 different labels (27 labels for the grid domains and one label for the boundary, see Supplementary Figure 10b). The relative difference between the local and global DA was defined as:

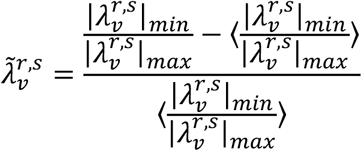

Here, 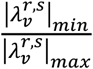 defines the ratio of the minimal to the maximal eigenvalue of the tensor 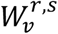, while 〈·〉 refers to the average value.

For the calculation of the quadratic and cubic invariants of the irreducible Minkowski tensors (IMT), the rank s was in the range of [0,8].

### 4.8. Ellipsoid representation of tensors

The rank-2 Minkowski and MIL tensors were visualized by ellipsoid surfaces, with radii and directions that were defined by the eigenvalues and eigenvectors of the tensors, respectively [38, 41]. The surface of an arbitrarily oriented ellipsoid, centered at the origin is obtained by solving the following expression for ***x***:

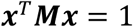

where ***M*** is a positive-definite matrix. The eigenvalues *λ_s_* of ***M*** are related to the semi-axes *c_s_* of the ellipsoid by:

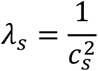

The corresponding eigenvectors then represent the principal orientations of the ellipsoid. By exploiting this relationship, the eigenvalues and eigenvectors of the rank-2 tensors used in this work could be represented as ellipsoid surfaces. In visualizing the ellipsoids of the spatially decomposed samples, the ellipsoids were uniformly scaled to the same volume, which is given by:

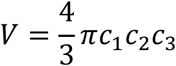

### 4.9. Standard bone morphometric analyses

The mean intercept length (MIL) analysis was performed by applying the algorithms implemented in BoneJ (version BoneJ2), which is an ImageJ plug-in, to the binary image stacks of the trabecular specimens [42]. While the earlier implementations of the MIL could suffer from significant deviations in the predicted anisotropy due to sampling bias [38, 43], the current implementation draws test lines through the entire image stack, offering a more uniform sampling. A convergence analysis was performed to assess the influence of the number of the parallel test lines and the number of different test line directions. For the final analyses, 2000 directions and 10000 lines per direction were used. Moreover, the MIL results for every sample were taken as the average of three runs of the MIL algorithm. The Euler characteristic (*χ*) was also estimated using BoneJ, on the same binary image data as was used for the MIL analysis. Since the estimation of the Euler characteristic assumes a single connected component, the images were purified (using the Purify command in BoneJ) prior to the connectivity computation. The other reported morphometric indices (*i.e*., BS and *∂S/∂r*) were obtained from the Scanco micro-CT scanner software (Scanco Medical AG, Switzerland).

### 4.10. Statistical analysis

The Kruskal-Wallis H test was used to detect significant differences between the means of the different groups of data. *Post hoc* comparisons of the means were performed using the two-sided Mann-Whitney U tests. The two-sample Kolmogorov-Smirnov test was used to assess the differences between the probability distributions of the *q_s_* and *w_s_* metrics, and the age-related curvature probability distributions. The obtained results were considered to be statistically significant when *p* < 0.01. All statistical analyses were performed using the python library Scipy.

